# Building the *lac* operon: A guided-inquiry activity using 3D-printed models

**DOI:** 10.1101/2020.01.14.904847

**Authors:** Claire L. Gordy, Conner I. Sandefur, Tessa Lacara, Felix R. Harris, Melissa V. Ramirez

## Abstract

Undergraduate biology courses rely heavily on visual representation of information. Students view images of plants, animals, and microbes, interpret data presented in graphs, and use drawings to understand how cells and molecules interact in three dimensions. Traditional teaching approaches exclude students with visual impairments and disadvantage students with disabilities that affect their interpretation and processing of visual and spatial information as well as students who simply do not identify as “visual learners”. By using new technologies to develop tactile teaching tools (TTTs) that can be employed in classrooms, we aim to create inclusive learning environments and more effectively instruct diverse learners. The advent of affordable and accessible 3D printing technology makes it possible to create tactile models that represent molecules, cells, and entire organisms more accurately than traditional visual representations. We describe the assessment of a 3D gene expression puzzle as a guided inquiry learning activity in which students must correctly assemble a series of components in order to achieve an output. Upon completion of the puzzle, the TTT provides tactile feedback through vibration to signal transcriptional activation. Analysis of pre- and post-assessment performance demonstrated statistically significant increases in individual students’ paired assessment scores in two different classroom implementations, with a greater effect size at a rural minority-serving institution than an urban R1 university. These encouraging preliminary data suggest that TTTs with guided-inquiry learning disproportionately benefit disadvantaged student populations and could serve as a tool in leveling the playing field when teaching abstract biological concepts in diverse educational settings.

## INTRODUCTION

Recent advances in 3D printing have led to the development of 3D printing labs termed “Makerspaces” at colleges and universities across the nation (1). Through these Makerspaces, instructors have begun to incorporate 3D-printed cellular and molecular models into their courses. Many of these 3D models serve solely to accurately represent a structure in three dimensions. While visualizing biology in three dimensions is critically important, simply replacing two-dimensional images with three-dimensional models without altering the way the information is presented represents a passive approach to learning. Furthermore, passive presentation of either two-dimensional or threedimensional images fails to include blind students and students with disabilities affecting their vision or processing of visual information.

We have developed an alternate strategy in which students utilize 3D-printed Tactile Teaching Tools (TTTs) paired with constructivist classroom activities to actively build an understanding of biological processes (Ramirez and Gordy, manuscript submitted). TTTs are 3D models composed of multiple parts representing interacting components, such as proteins of a signaling pathway or monomers that compose a biological macromolecule. These models are used for guided inquiry learning (GIL) activities that ask students to manipulate, assemble, or analyze structures in order to answer questions. Based on the structure of Process-Oriented Guided Inquiry Learning (POGIL) activities (2, 3), the GIL activities paired with TTTs require students to work in groups. Each group begins by exploring and defining the individual pieces of the TTT, and questions gradually increase in complexity, with each step building on the previous. In contrast to existing POGIL activities, TTTs with GIL activities allow an additional level of constructivist learning: students can experiment with the pieces of the TTT to predict and test outcomes.

Here we describe a TTT and GIL activity in which students work in small groups using 3D interactive models to define the structure of the *lac* operon and construct an understanding of basic mechanisms of gene regulation. Both the *Vision and Change in Undergraduate Biology Education* report and the American Society for Microbiology (ASM) curriculum guidelines have identified gene expression as a core concept for undergraduate biology students (4, 5). The *lac* operon is commonly used to teach key principles related to gene organization and regulation of gene expression in undergraduate courses ranging from introductory biology to microbiology and genetics, yet this topic can be challenging for students (6, 7). Because this particular biological example is so widely utilized to illustrate the process of gene expression, the TTT and GIL activity described here focuses on the *lac* operon.

This TTT and GIL activity were designed to represent the interactions that regulate *lac* operon expression in three dimensions and to facilitate constructivist learning. The *lac* operon TTT utilizes magnets to simulate the binding of the LacI repressor to the operator and a vibration motor housed in the DNA model to indicate successful binding of the RNA polymerase to the relevant promoter sequences and initiation of transcription. As they are guided through their exploration of the TTT, students first construct an understanding of the individual components and then how the components interact to further build their understanding of the biological process of gene expression. Finally, students are asked to apply their understanding in new contexts by predicting outcomes based on particular interactions.

While numerous studies have shown the benefits of active learning over traditional lecture methods, more recent studies have demonstrated that these strategies are more impactful for certain underrepresented minority (URM) groups (8–11). We hypothesized that the use of the *lac* operon TTT-GIL would result in increased student performance on assessments of related learning objectives in multiple educational settings, and that learning gains would be greater for students at a less resourced minority-serving institution (MSI) than at an R1 university.

### Intended audience and prerequisite knowledge

This activity is intended for an introductory microbiology or genetics course for life sciences majors. It could be simplified for use in a general biology course. Students should have a working understanding of the central dogma of molecular biology.

### Learning time

This learning activity was implemented using 75 minutes of classroom time. The activity can be completed in a single class session or spread over multiple class sessions depending on class schedule and format.

### Learning objectives

At the end of this activity, students should be able to:

1. Diagram a bacterial operon including the promoter, −10/−35 sites, and transcriptional and translational start/stop sites.
2. Describe how the *lac* operon is regulated.
3. Explain how *lac* operon regulation enables a bacterium to effectively utilize two different sugars as growth substrates under catabolite repression control.

## PROCEDURE

### Materials

Each student group is provided with a bag containing all of the *lac* operon components (DNA, repressor protein, allolactose, and RNA polymerase) (Figure 1). The required materials and detailed instructions for their creation can be found in Appendix 1.

**Figure 1:**
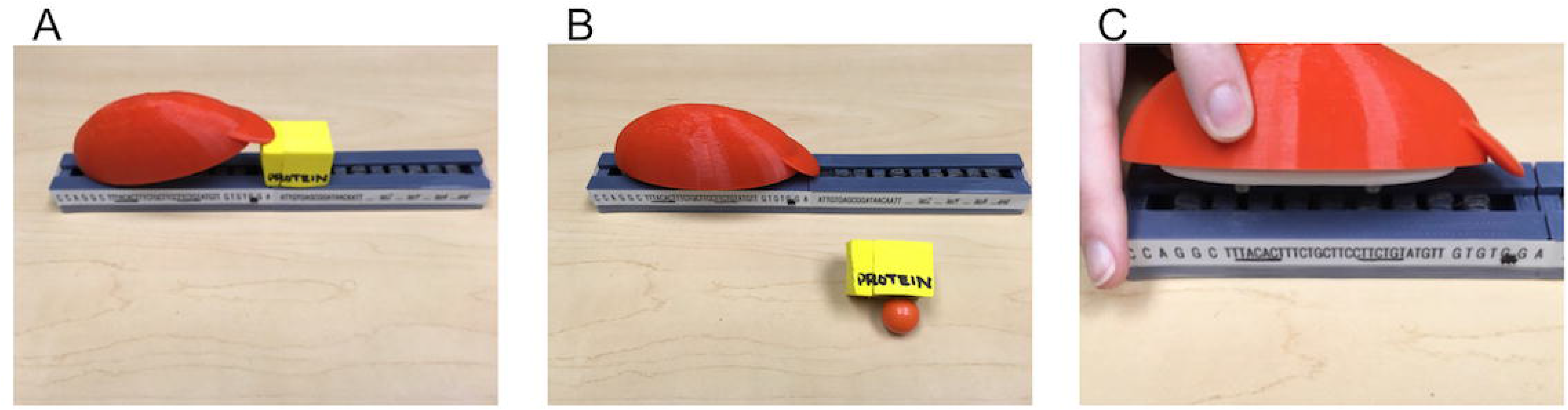
3D-printed *lac* operon tactile teaching tool set. **A**. the yellow LacI repressor protein is shown binding to the operator site within the promoter. **B**. The allolactose (magnetic marble) is shown binding the LacI repressor, pulling it off of the operator sequence. With LacI unbound, the red/white RNA polymerase is able to bind to specific −10/−35 regions in the *lac* promoter as shown in **C**. Magnets are used to simulate interactions between components. The model utilizes a simple circuit with the battery contained within the RNA polymerase and a vibration motor contained in the gray DNA box. When the RNA polymerase binds to specific −10/−35 regions in the *lac* promoter, the circuit is completed, and the model begins to vibrate, representing transcriptional initiation.

### Student instructions

The student GIL worksheet packet is provided in Appendix 2, with an instructor key in Appendix 4. Ideally, students should work in small groups of approximately three to ensure that all students have the opportunity to manipulate the TTTs. Each student should complete the GIL worksheet packet individually, but students are encouraged to talk through the activity in their groups. The GIL activity is broken into seven parts. Students should pause after each section and not work ahead. Instructions for directing student interactions with the TTT are provided in the GIL worksheet packet.

### Faculty instructions

#### Preparation of TTTs

3D print files and step-by-step instructions for assembly and wiring are included in Appendix 1 and posted online at https://stembuild.ncsu.edu/resource/microbial-genetics-the-lac-operon/.

#### Pre-activity lecture

Prior to the TTT and GIL activity, students should review or be introduced to the processes captured in the central dogma of molecular biology. The amount of classroom time dedicated to this introduction will vary depending upon the students’ prior experience with this material, and this introductory lecture could occur in a separate class period before the activity. After this introductory lecture, a modified case study entitled “Cow of the Future: Genetically Engineering a Microbe to Reduce Bovine Methane Emissions” (12) can be used to introduce the basic principles of microbial genetics. We have implemented the TTT and GIL activity both with and without the case study, but in both cases, used the questions associated with the case study as a pre-assessment.

#### TTT-GIL Intervention

During the class period when the TTT-GIL activity is implemented, faculty should have students break into small groups to complete the GIL worksheet packet. In both implementations of this activity, the worksheet packet (Appendix 2) was supplemented with published POGIL activities covering transcription and translation (13, 14). Depending on the needs of the class, instructors may wish to also include a published POGIL activity covering bacterial gene regulation (15). After all groups complete each section, the instructor should lead a classroom discussion reviewing what the students have just completed.

### Suggestions for assessing student learning

Possible summative assessment data that could be obtained include student performance on pre-assessment questions from the “Cow of the Future: Genetically Engineering a Microbe to Reduce Bovine Methane Emissions” case study available from the National Center for Case Study Teaching in Science (12), student performance on the GIL worksheet packet (Appendix 2), or student performance on post-assessment questions (Appendix 3). Possible formative assessment strategies include muddiest point activities done at the end of the class period in which the TTT-GIL activity was implemented. At the start of the next class period, the instructor could discuss any questions that emerged.

### Safety issues

For the printing of the *lac* operon, the safety guidelines of the specific 3D printer being used should be followed. Possible risks associated with 3D printing include heat, electrical, mechanical, and fume risks. There are no risks associated with the use of the assembled models in the classroom setting.

## DISCUSSION

### Field Testing

This activity was implemented in two separate courses at two different institutions: North Carolina State University (NCSU), an urban public R1 university, and the University of North Carolina - Pembroke (UNCP), a rural public minority-serving institution (MSI) (Table 1). The first iteration of this activity occurred in a 300-level introductory microbiology course at NCSU in which 57 students were assessed. Of note, this section of the course was taught in the SCALE-UP format (Table 2). SCALE-UP (Student Centered Active Learning Environment with Upside-down Pedagogies) is a type of learning environment that encourages interaction by utilizing both design and technology (16, 17). In the SCALE-UP classrooms at NCSU, students sit together at round tables of nine students, and frequently work in groups of between three and nine students. The course content can be delivered using both lecture and in-class group work that promotes active learning.

**Table 1:**
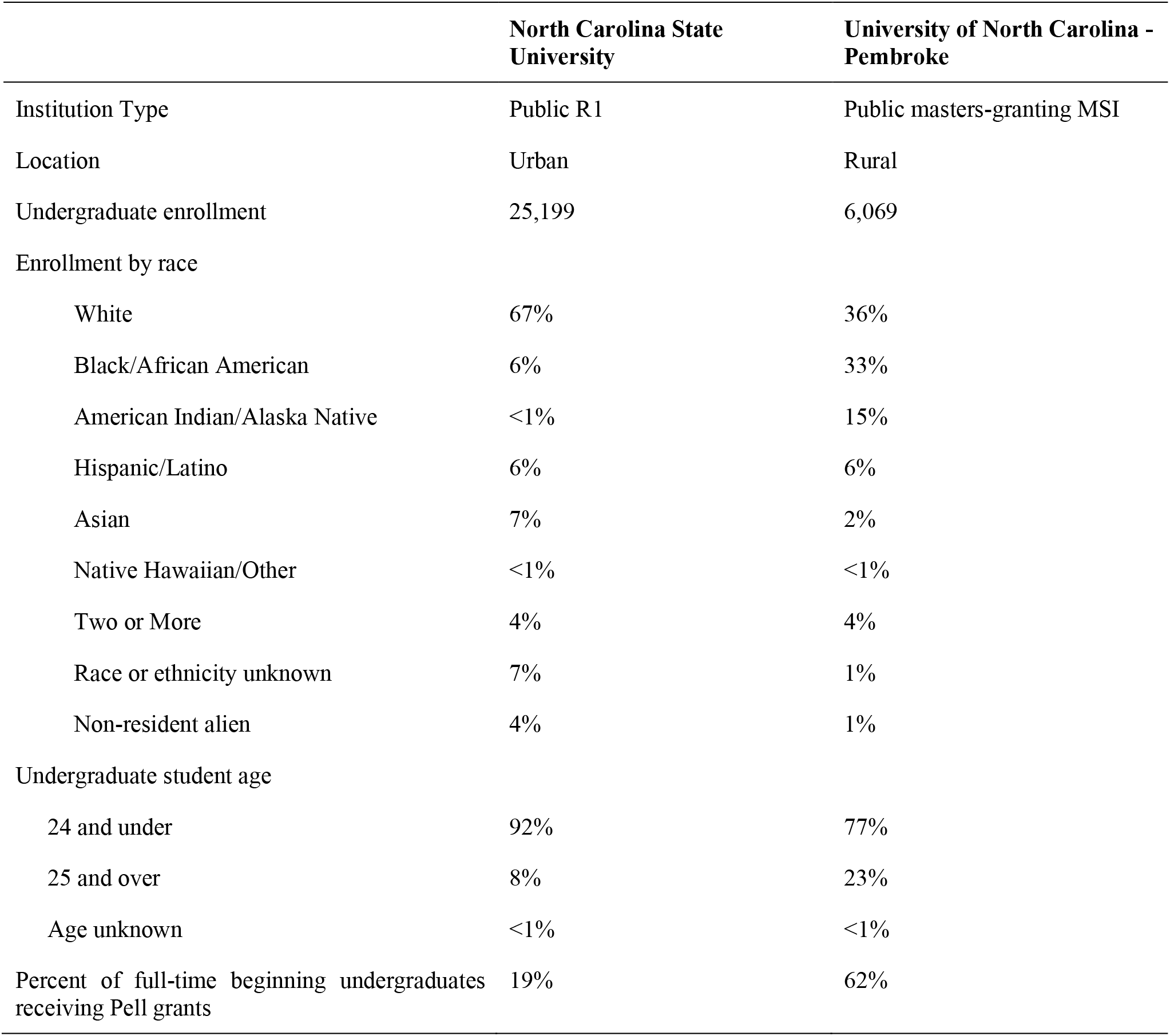
Institutional characteristics.

**Table 2:** Characteristics of classrooms in which *lac* operon TTT-GIL activity was implemented and assessed.

| **Institution** | **Course implemented** | **Enrollment (Participants)** | **Iteration** |
| --- | --- | --- | --- |
| NCSU | 300-level microbiology | 58 (57) | 1 |
| UNCP | 300-level genetics | 44 (29) | 2 |

In this microbiology course at NCSU, the section on microbial genetics spans approximately four class periods. The pre-assessment (Appendix 3) was given during the first class period of the section following a 45-minute lecture focused on the central dogma of molecular biology. The following two class periods continued to build on the principles of microbial genetics by introducing horizontal gene transfer and microbial gene expression. In this semester, the fourth class period was devoted to the *lac* operon TTT and GIL activity (intervention). Working in groups, students were given a *lac* operon kit with prompts directing them how to interact with their kit in order to answer a series of questions related to the central dogma and microbial gene expression.

The activity begins with questions from published POGIL activities (13, 14) and increases in complexity as students move from studying two-dimensional images to the three-dimensional TTT. Upon assembly of the puzzle, the TTT provides tactile feedback through vibration to signal transcriptional activation. Based on the locations at which the polymerase model interacts with the DNA model to initiate vibration, the students label the locations of the −35, −10, and +1 sites. As the activity progresses, the students add a LacI repressor protein, which interacts with the DNA via a magnet to prevent RNA polymerase binding, and a magnetic marble representing allolactose, which binds LacI to prevent its interaction with the DNA. By experimenting with the parts of the model to uncover these interactions, the students are able to identify the location of the *lac* operator and predict how changes in lactose levels will affect transcription of the genes encoded in the operon.

The subsequent exam included questions similar to those used in the pre-assessment, but using an alternative operon example (post-assessment, Appendix 3, Table 3).

**Table 3:** Pre- and post-assessment questions aligned with learning objectives.

| Question | Pre-assessment: | Post-assessment: | Learning objective |
| --- | --- | --- | --- |
| 1 | How many mRNA molecules will be produced when this operon is transcribed? | How many mRNAs would be transcribed from this operon? | Diagram a bacterial operon including the promoter, -10/-35 sites, and transcriptional and translational start/stop sites. |
| 2 | How many promoters would be necessary to allow transcription of the *pmoCAB* operon? | How many promoters control expression of this operon? |  |
| 3 | How many start codons and stop codons will be present on the RNA produced when the *pmoCAB* operon is transcribed?* | How many start codons would be present in this operon? |  |
| 4 |  | How many stop codons would be present in this operon? |  |

\* Student responses to question 3 were scored separately for the start and stop codons (for example, a student who selected “3 start, 1 stop” would receive credit for start codons but not for stop codons) to allow matching with the post assessment, in which two separate questions about start and stop codons were included (3 and 4).

In an effort to evaluate the impact of this approach at different types of institutions, a collaboration was established with faculty at UNCP. UNCP is an MSI with a historical mission of serving the American Indian population of rural southeastern North Carolina. Located in Robeson County, one of the poorest counties in the US (18, 19), UNCP has a significantly higher proportion of students receiving Pell grants than NCSU (62% vs. 19%; Table 1) (20). The activity was implemented in one 300-level introductory genetics course, and 29 students were assessed.

The activity was originally scheduled for the second unit of the semester, which usually covers expression of genetic information, including two weeks focused on transcription and translation. This material is typically taught through two POGIL activities and two lectures with clicker questions (over the course of a week) on regulation of gene expression in eukaryotes and prokaryotes. Due to Hurricane Florence, UNCP was closed for approximately two weeks. In order to accommodate prior scheduling of guest speakers for the course, the regulation of prokaryotic gene expression was moved to the end of the semester, which was extended one week. Prior to beginning the activity, students had a 20-minute lecture with clicker questions reviewing the central dogma. Afterwards, students completed the pre-assessment (Appendix 3). The activity was carried out over 1.5 50-minute class periods. Working in groups of three or four, students worked on the activity in the second half of the first class period and completed the activity during the second class period. The post-assessment (Appendix 3) was given the next week during the final exam period.

### Evidence of student learning

De-identified, linked performance data was compiled from pre-assessments performed before the TTT-GIL intervention and post-assessment questions included on the unit exam, allowing analysis of both population-wide trends and individual students’ learning gains (Table 3). Analysis of linked pre- and post-assessment scores demonstrated that the NCSU and UNCP interventions both resulted in statistically significant increases in performance on an individual level as well as increases in median assessment scores (NCSU median pre-assessment: 75% vs. median post-assessment: 100%; UNCP median pre-assessment: 25% vs. median post-assessment: 75%; Fig. 2A). Both populations demonstrated similar normalized average learning gains (Hake’s *g*, calculated as 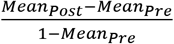 (Table 4) (21). Because Hake’s *g* has been shown to be biased in favor of populations with high pre-assessment scores (e.g., the NCSU population), we also measured effect size (Cohen’s *d* with correction for small sample size, calculated as 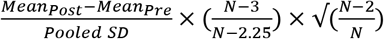, where Pooled 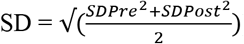) (22, 23). Strikingly, the effect size in the UNCP population was 0.82, while the effect size in the NCSU population was 0.50. Together, these results demonstrate that while both groups of students benefitted from the TTT-GIL activity, the effect of this activity was greater for students at UNCP.

**Figure 2:**
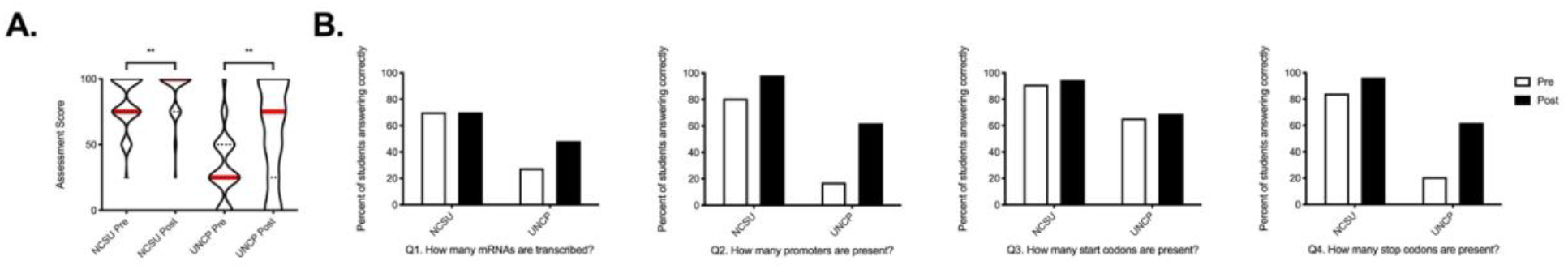
Impact of *lac* operon guided-inquiry learning activity on student learning demonstrated by performance on pre- and post-assessments. **A**. Pre- and post-assessment scores are shown as violin plots. The width of the plot represents the percent of students receiving a given score. Dotted lines represent quartiles, and solid red lines represent medians. Individual students’ linked pre- and post-assessment scores were measured using a Wilcoxon matched pairs signed-rank test (****, p < 0.0001,) **B**. The percent of students answering each assessment question correctly is shown for NCSU (left) and UNCP (right). Open bars represent the pre-assessment, and filled bars represent the post-assessment.

**Table 4:** Normalized learning gains and effect sizes.

| **Institution** | **NCSU** | **UNCP** |
| --- | --- | --- |
| Mean Pre-assessment Score | 81.58% | 32.76% |
| Pre-assessment SD | 19.21% | 25.09% |
| Mean Post-assessment Score | 90.79% | 60.34% |
| Post-assessment SD | 16.11% | 36.91% |
| Pooled SD | 17.73% | 31.56% |
| Sample Size | 57 | 29 |
| Average Gain | 9.21% | 27.59% |
| Maximum Possible Gain | 18.42% | 67.24% |
| Hake's $g$ | 0.50 | 0.41 |
| Cohen's $d$ | 0.50 | 0.82 |

Furthermore, examination of student performance on individual assessment questions reveals some interesting trends (Figure 2B, Table 5). On the pre-assessment, both groups performed best on question 3 (How many start codons are present?), and both groups showed small improvements on the post-assessment (NCSU: 91.2% correct answers pre vs. 94.7% post; UNCP: 65.5% pre vs. 69.0% post). On questions 2 and 4 (How many promoters are present? and How many stop codons are present?, respectively), NCSU students performed at a high level on the preassessment (80.7% and 84.2% correct answers, respectively), and improved to 98.2% and 96.5% correct answers, respectively. In contrast, UNCP students performed at a low level on these questions on the pre-assessment (17.2% and 20.7% correct answers, respectively), but improved their performance to 62.1% correct answers on both questions. Interestingly, on the question for which NCSU students had the most room for improvement -- question 1 (How many mRNAs are produced?) – they showed no improvement (70.2% correct answers pre and post), while UNCP students improved from 27.6% to 48.3% correct answers.

**Table 5:** Percent of students answering correctly for each assessment question.

| Question | NCSU Pre | NCSU Post | UNCP Pre | UNCP Post |
| --- | --- | --- | --- | --- |
| 1 | 70.2% | 70.2% | 27.6% | 48.3% |
| 2 | 80.7% | 98.2% | 17.2% | 62.1% |
| 3 | 91.2% | 94.7% | 65.5% | 69.0% |
| 4 | 84.2% | 96.5% | 20.7% | 62.1% |

While the study design does not allow us to definitively determine the reasons why this activity benefitted UNCP students more strongly than NCSU students, this finding is in line with many studies demonstrating that increased structure and active learning strategies disproportionately benefit students from underrepresented or disadvantaged groups (8–11). Furthermore, differences in the undergraduate biology curricula between the two institutions may impact students’ baseline knowledge of concepts related to genomic organization and gene regulation prior to the intervention, as students at NCSU are likely to have encountered the *lac* operon and the central dogma of molecular biology in multiple courses prior to enrollment in general microbiology, while UNCP do not encounter the *lac* operon or the central dogma of molecular biology in any courses other than the genetics course in which this TTT-GIL activity was implemented. To determine whether this activity -- and TTTs in general -- specifically benefit underrepresented students and/or students with less background knowledge will require studies in which student demographic data and course enrollment history are collected or the intervention is performed in a lower-level course, such as introductory biology.

Together, these preliminary data indicate the utility of implementing TTTs with guided-inquiry learning as a strategy to enhance students understanding of abstract biological concepts, and suggest future avenues of exploring the effectiveness of this teaching strategy in leveling the playing field for students who belong to underrepresented groups or attend underresourced institutions.

## CONCLUSION

As Makerspaces become more commonplace and accessible to instructors and students, there is an opportunity to utilize emerging maker technologies for educational purposes. While 3D models have been utilized in the classroom in a variety of ways (Ramirez and Gordy, manuscript submitted), best practices for utilizing models in inclusive ways remain undefined. Based on these encouraging preliminary results, our group has expanded this approach to additional biological concepts covered in additional biological sciences undergraduate courses in a variety of classroom settings. Future studies continuing to compare the effectiveness of TTTs for different student populations will help to establish TTT-GIL activities as a new tool for inclusive STEM teaching.

## ACKNOWLEDGEMENTS

We would like to thank the NC State University Distance Education and Learning Technology Applications (DELTA), Dr. Robert Beichner, Director of the STEM Education Initiative, and the College of Sciences Faculty Research and Professional Development Program for financial support of this work. Additionally, we are grateful for the collaborative spirit of the NC State Libraries Makerspace staff and thank them for their expertise, support, and enthusiasm. We would also like to recognize our students for their willing participation. The authors declare that there are no conflicts of interest.

## SUPPLEMENTAL MATERIALS

Appendix 1: Instructions for printing and assembling the *lac* operon model

Appendix 2: Student worksheet with instructor notes

Appendix 3: Pre- and post-assessment questions

Appendix 4: Student worksheet key

Appendix 5: Pre- and post-assessment key

Appendix 6: Blank student worksheet

IRB Information - This study was performed following a protocol that was first exempt, and then later approved by the NC State University Institutional Review Board for the Use of Human Subjects in Research (Protocol Number 12731).

