## Appendices for "Building the *lac* operon: A guided-inquiry activity using 3D-printed models"

**Appendix 1: *lac* operon model printing and assembly instructions**

**3D Print Files are available for download here: [https://stembuild.ncsu.edu/resource/microbial-genetics-the-lac-operon/](https://stembuild.ncsu.edu/resource/microbial-genetics-the-lac-operon/)**
**Unless noted, the materials listed below are available at educational supply stores or Amazon.com.**

**MATERIALS NEEDED**

- Printer: Lulzbot Mini or similar
- Filament: PLA 2.85mm or appropriate filament for your printer/nozzle. You may wish to use three different colors (one for both parts of the DNA operon box - we used grey, one for the RNA polymerase top - we used red, and one for the RNA polymerase bottom - we used white).
- Items to 3D print
    - 2 x Top of DNA box
    - 2 x Bottom of DNA box
    - RNA Polymerase top
    - RNA Polymerase bottom
- 10mm Mini Vibration Motor with wire leads or light (1 per model)
- 10mm ceramic disc magnets (2 per model)
- CR2032 3-volt battery and its respective holder (1 per model)
- Thin conductive wire for lead connection
- 4-40 x 1/2 Socket Head Cap Screw (McMaster 92196A110) (2 per model)
- Magnetic marbles ⅝" diameter (1 per model)
- 1" foam cubes (2 per model)
- Conductive thread (~12 inches per model)
- Hot glue gun
- Wire clippers
- Needle nose pliers
- Label maker and adhesive label tape
- Sharpies

**PRINTING INSTRUCTIONS**

- Collect the materials necessary for 3D printing:
    - Filament
    - 3D printer
    - Laptop

- Follow the standard procedures for your Makerspace/your printer to print each of the four parts of the model: the RNA polymerase top and bottom, the DNA operon box top and bottom. We recommend printing the box top upside down (inside facing upwards) with supports so any elephant foot effect that happens doesn't interfere with the fit of the boxes. The bottom boxes should be printed bottom down for this same reason.

- Suggested settings:
    - Infill: 20%
    - Supports: 15%

**ASSEMBLY INSTRUCTIONS: RNA polymerase**

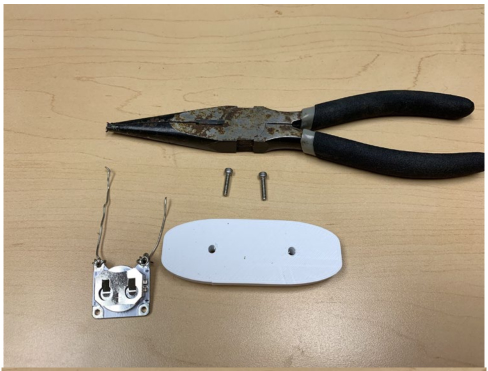

- Print RNA polymerase top and bottom as described in print instructions.
- Collect RNA polymerase supplies needed - battery, battery holder, thin conductive wire, screws, needle nose pliers.

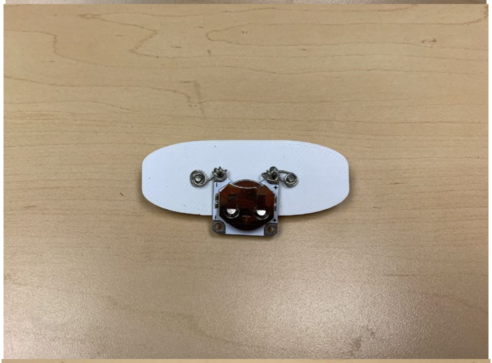

- Wrap wire around the battery leads, and place the battery in the holder. Alternatively, battery holders with wire leads already attached can also be purchased.

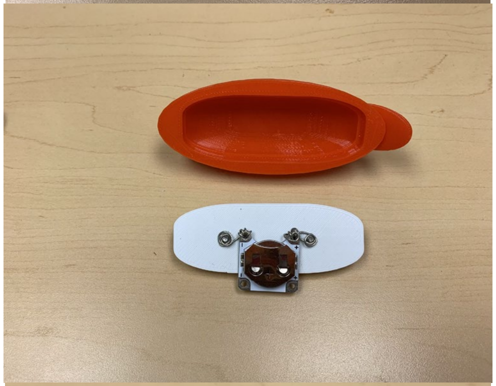

- Place the screws through the plastic plate from the rounded side through to the flat side. Wrap the wires around the length of the screw to secure the battery holder to the flat side of the plate.

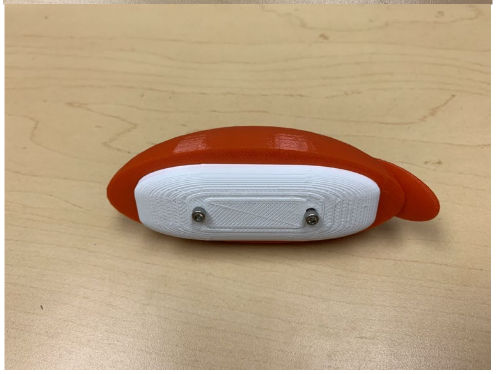

- Pop the plastic plate onto the bottom on the RNA polymerase top.

**ASSEMBLY INSTRUCTIONS: *lac* operon boxes**

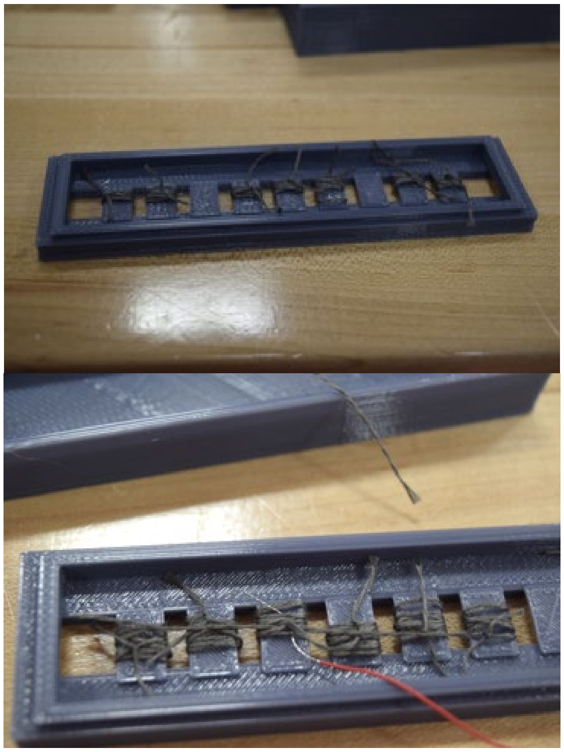

*lac* operon promoter region sequence:

CCAGGCTTTACACTTTCTGCTTCCTTCGTTA
TGTTGTCGG**A**ATTGTGAGCGGATAACAATT

*Notes: -35/-10 sequences are underlined, +1 is bolded in red*

DNA box 1 - *lac* promoter region

- Print DNA operon box top and bottom as described in print instructions.
- Place the operon box top face down, as shown in the image. Note that the box top has 9 tiles.
- Obtain about 6 inches of conductive thread and wrap thread around each individual tile on the DNA top. After wrapping each tile with thread, tie off with a knot and cut off any excess thread. Note: make sure you thread on the bottom of the DNA top so thread strands are not exposed to the top track.
- Tiles 3 and 7 are the contact points for the RNA polymerase. The conductive thread wrapped around these tiles will contact the vibration motor wire.
- Strip the ends of the vibration motor wire leads using wire clippers.
- Insert the stripped red wire into the wrapped conductive thread on tile 3, and the stripped black wire into the wrapped conductive thread on tile 7.
- Hot glue magnet onto the furthest tile from the black wire.
- Snap pieces together and test by running RNA polymerase along the track. The motor should vibrate when the RNA polymerase contacts tiles 3 and 9. If it does not, check and adjust wiring as necessary.
- Print the *lac* promoter region sequence onto a label and adhere to the front of the DNA box. Format spacing appropriately to the size of the DNA box.
- For accessible *lac* promoter sequence, we have utilized the website www.touchesee.me to convert the sequence to braille and export as .stl files for 3D printing. These can be adhered to the front of the DNA box.

*lac* operon gene region:

*lacZ* *lacY* *lacA*

DNA box 2 - *lac* gene region

- Obtain about 6 inches of conductive thread and wrap thread around each individual tile on the DNA top. After wrapping each tile with thread, tie off with a knot and cut off any excess thread. Note: make sure you thread on the bottom of the DNA top so thread strands are not exposed to the top track.

- Snap the bottom and top pieces together.
- Print the *lac* gene region onto a label and adhere to the front of the DNA box. Format spacing appropriately to the size of the DNA box.
- For accessible *lac* gene labels, we have utilized the website [www.touchesee.me](http://www.touchesee.me) to convert the sequence to braille and export as. stl files for 3D printing. These can be adhered to the front of the DNA box.

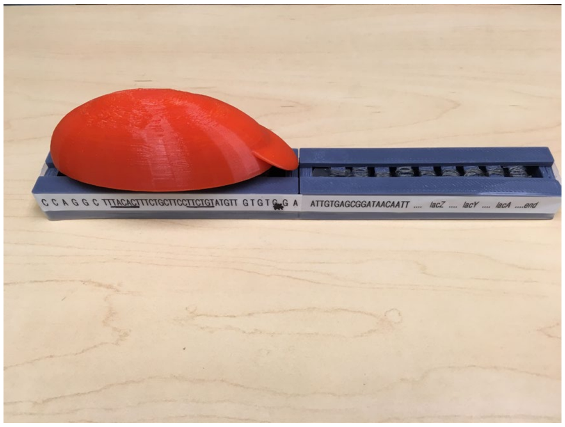


- Glue the two DNA boxes together with the promoter box to the left of the gene box.

**ADDITIONAL**

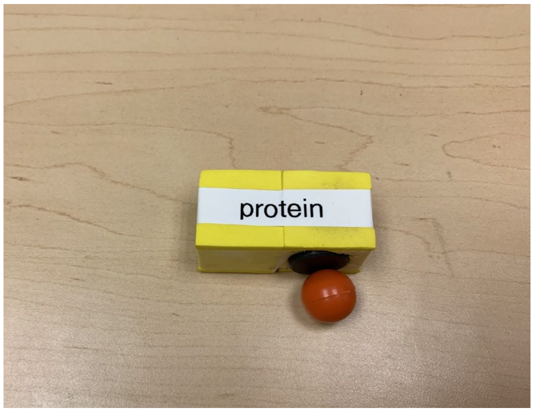


- Glue two 1” foam cubes together to make the repressor protein. Print a label for the protein. Glue a magnet to the bottom right side of the protein. The foam can easily be cut away to allow the magnet to be depressed inside of the cube. Before gluing down make sure the magnet orientation is correct for binding to the magnet already placed inside the DNA box described above.. This is where the magnetic marble (allolactose) will bind.

**Appendix 2: Student worksheet with instructor notes**

### PART 1: MOLECULAR BIOLOGY BASICS

*Instructor Note: PART 1 is modified from the activity "POGIL Gene Expression – Transcription", POGIL™ Activities for AP* Biology. We recommend using whatever portions of the activity are appropriate for your course in providing a review of the central dogma.*

### PART 2: BACTERIAL GENOMIC ORGANIZATION

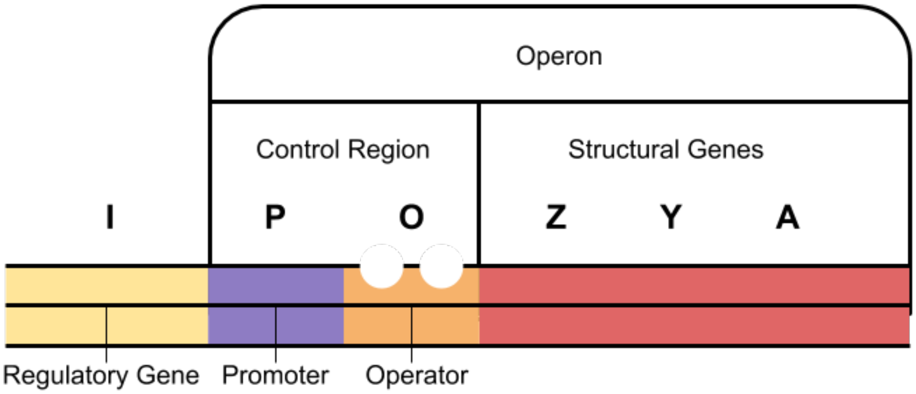


1. Refer to the image above to answer the following questions.

   a. How many genes are contained in this operon?

   b. How many promoters control the expression of this operon?

   c. How many mRNAs would be transcribed from this operon?

   d. How many polypeptides would be produced?

   e. How many start codons, and how many stop codons would be present in this operon?

   f. How many ribosome binding sites (RBS) would be present in this operon?

### PART 3: BACTERIAL TRANSCRIPTION

1. Where in the eukaryotic cell does transcription take place?
2. Where in the prokaryotic the cell does transcription take place?
3. List the components that would be included in a bacterial **promoter** region.

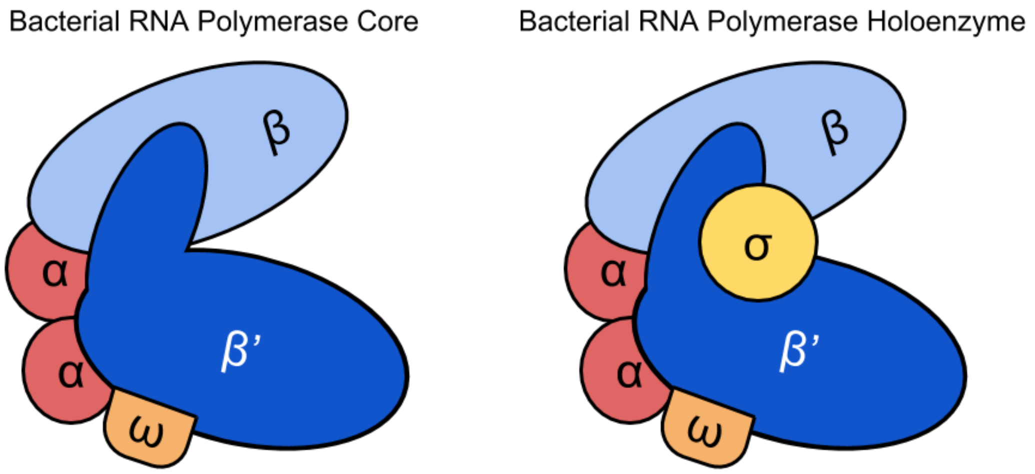


4. The image above shows the bacterial RNA polymerase core and holoenzyme complexes. What additional subunit is present in the holoenzyme that is not part of the core?

5. Which subunit(s) do you think would bind each of the promoter regions that you listed in Q3?

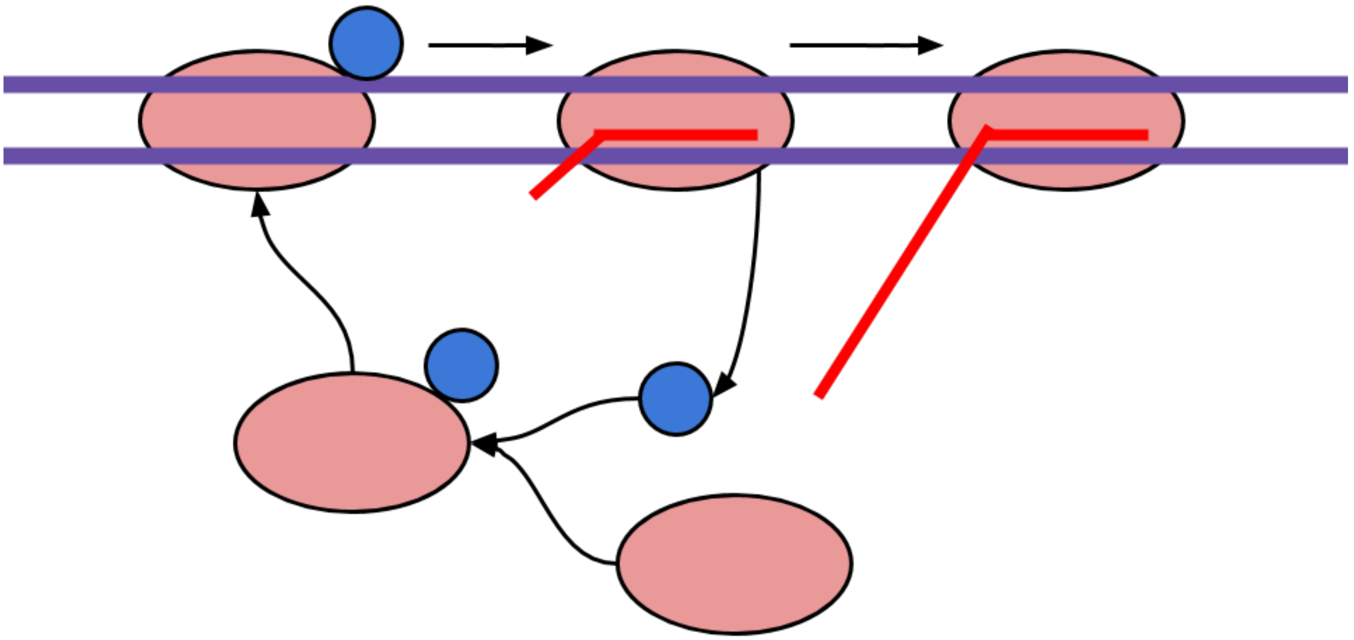

6. The image above shows a cycle that occurs during transcription initiation and elongation.

   a. What do you think the blue ball represents?

   b. What do you think the red oval represents?

   c. What do you think the red line represents?

d. Describe what is happening in this image.

#### PART 4: BACTERIAL TRANSLATION

*Instructor Note: PART 4 is modified from the activity "POGIL Gene Expression – Translation", POGIL™ Activities for AP* Biology. We recommend using whatever portions of the activity are appropriate for your course in providing a review of translation.*

#### STOP

##### COMPLETE PART 5 WITH THE MODEL PROVIDED NOW.

#### PART 5: REGULATION OF BACTERIAL GENE EXPRESSION

1. Examine the red/white model.
   a. What do you think the red portion represents?

   b. What do you think the white portion represents?

2. Examine the gray DNA operon model. *Note- this model is only showing you the top strand (5'>3') for simplicity.

   a. How many promoters control the expression of this operon?

   b. Place the "feet" of the red/white model into the track on the left most side of the DNA. Slowly slide it towards the right, along the sequence until you arrive at the -35 and -10 regions.

   c. Underline the -35 and -10 consensus sequences using the black sharpie.

   d. Mark the approximate location +1 with an *

3. Examine the gray DNA operon model. *Note- this model is only showing you the top strand (5'>3') for simplicity.

   a. List the genes that are contained in this operon.

   b. How many mRNAs would be transcribed from this operon?

   c. How many polypeptides would be produced?

   d. With your black sharpie, mark the location where you would find the START and STOP codon(s).

e. With your other color sharpie, mark the location where you would find the RBS(s).

4. Using the foam "protein", determine where on your DNA operon model this protein will bind.

   a. Underline this sequence (does not need to be precise) with your color sharpie.

   b. What protein does the foam protein represent?

5. Allow the pink marble to "interact" with the foam protein.

   a. How does the pink molecule interacting with the foam protein impact the red/white model from performing its function?

   b. What molecule does the pink molecule represent?

   6. If the pink marble is not present, is the transcription of this operon turned on or off?

   7. If the pink marble is present, is the transcription of this operon turned on or off?

#### PART 6: REGULATION OF BACTERIAL GENE EXPRESSION

Recall that glucose is used to generate energy in the form of ATP through glycolysis, and in some organisms, through oxidative phosphorylation. It is the preferred carbon source for *E. coli*.

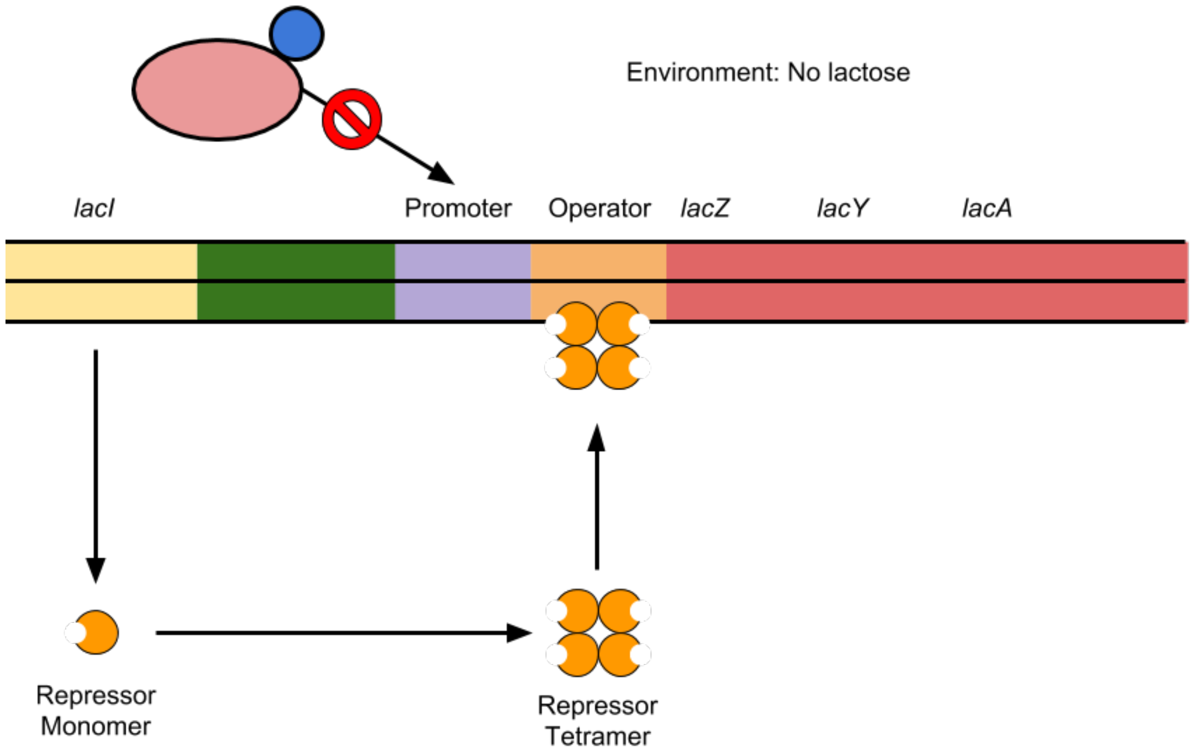


1. Refer to the image above to answer the following questions.

   a. What is encoded by the *lacI* gene?

   b. Is the *lacI* gene part of the lac operon?

   c. What happens to the protein encoded by the *lacI* gene in the absence of lactose?

   d. To what site in the DNA does the protein bind?

   e. How does this affect the transcription of the genes in the *lac* operon?

Environment: Lactose Present

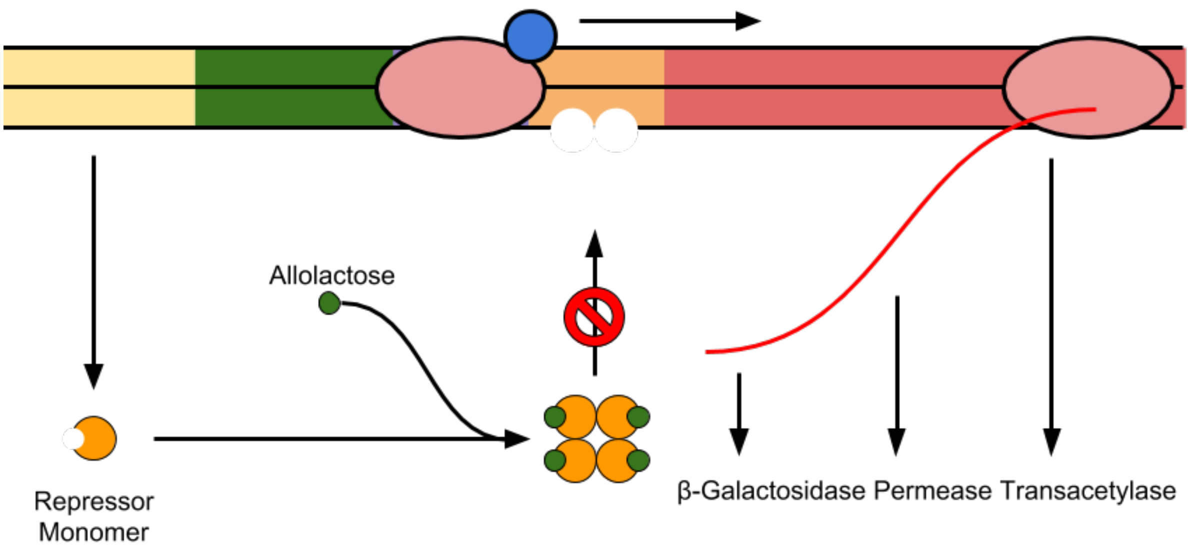


2. Refer to the image above to answer the following questions. Note, when lactose is present in large enough quantities, some of this lactose will be converted to allolactose.

   a. What molecule is considered the "inducer" in this system?

   b. What molecule or molecules are bound by the inducer?

   c. How does that affect the function of these molecules?

   d. How does this affect the transcription of the genes in the *lac* operon?

3. Considering both images together, how would you expect transcriptional activity at the *lac* operon to change if you added large quantities of lactose to an *E. coli* culture?

4. How would you expect transcriptional activity at the *lac* operon to change if you removed lactose from an *E. coli* culture?

5. Do you think that glucose levels should also regulate *lac* operon transcription? If so, how? That is, should high levels of glucose increase or decrease *lac* operon transcription?

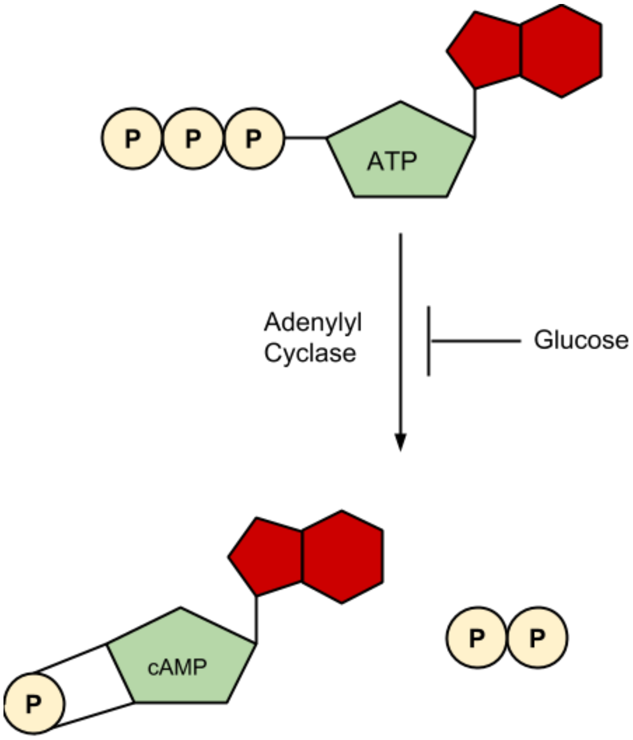


6. Describe the chemical reaction to the left in one or two sentences.

7. When glucose levels in the cell decrease, would you expect an increase or decrease in cAMP levels? Why?

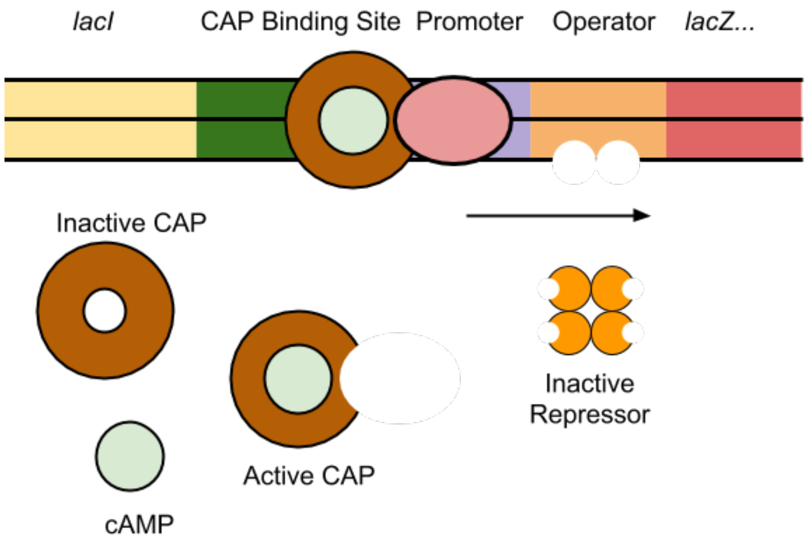


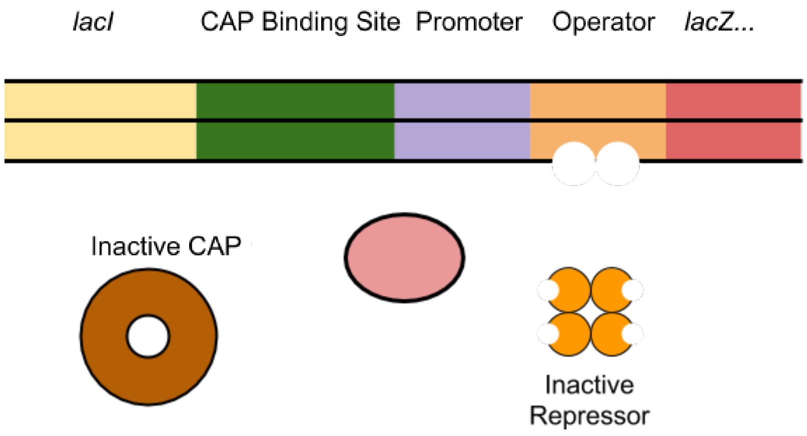


8. Refer to the image above to answer the following questions. In the images above, we have added some additional detail.

   In the top image, lactose is present, and glucose is absent.

   a. Where is the repressor? Why?

   b. Are cAMP levels high or low? Why?

   c. What protein is cAMP interacting with?

   d. What does this protein do when it interacts with cAMP?

e. How does this affect the transcription of the genes in the lac operon?

In the bottom image, lactose is present, and glucose is present.

f. Where is the repressor? Why?

g. Are cAMP levels high or low? Why?

h. What happens to CAP?

i. Are the genes in the *lac* operon transcribed?

#### PART 7: APPLYING THIS INFORMATION

1. What conditions are necessary for transcription of the *lac* operon to occur?

2. Would you say that lactose directly or indirectly regulates the *lac* operon?

3. Would you say that glucose directly or indirectly regulates the *lac* operon?

4. Come up with an example a mutation in one component of the *lac* operon, and answer the following questions.

   a. What component is mutated, and what is the mutation?

   b. In your altered version of the *lac* operon, the presence of lactose and absence of glucose, if the *lac* operon transcribed?

   c. Explain, or draw, why this is the case.

**Appendix 3: Pre and post assessment questions**

**Pre-assessment:** The pre-assessment consisted of questions adapted from the case study entitled "Cow of the Future Genetically Engineering a Microbe to Reduce Bovine Methane Emissions" (Stewart et al, 2014). Included with these questions was the pmoCAB operon diagram from the case study. Individual questions included as Question 3 in the published case study were formatted as multiple choice questions. Three of these questions were used as the pre-assessment for this activity:

1. How many mRNA molecules will be produced when this operon is transcribed?

2. How many promoters would be necessary to allow transcription of the pmoCAB operon?

3. How may start codons and stop codons will be present on the RNA produced when the pmo operon is transcribed?

Student responses to question 3 were scored separately for the start and stop codons (for example, a student who selected "3 start, 1 stop" would receive credit for start codons but not for stop codons) to allow matching with the post assessment, in which two separate questions about start and stop codons were included (3 and 4).

**Post-assessment:** The following exam questions were used for post-assessment.

The following is a diagram of the *ara* operon in *E. coli*. This operon encodes the enzymes needed to utilize the sugar arabinose.

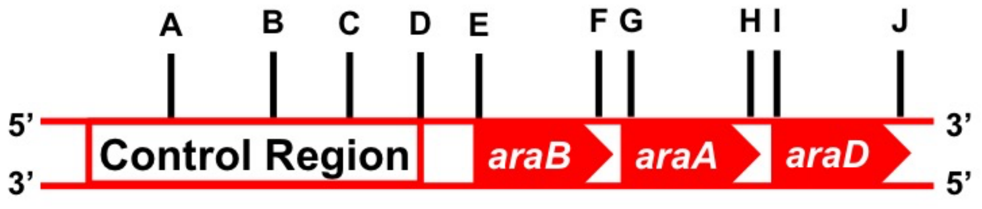


1. How many mRNAs would be transcribed from this operon?

Select one:

1

2

3

4

5

6

2. How many promoters control expression of this operon?

Select one:

1

2

3

4

5

6

3. How many start codons would be present in this operon?
Select one:

1

2

3

4

5

6

4. How many stop codons would be present in this operon?
Select one:

1

2

3

4

5

6

**Appendix 4: Student worksheet with instructor notes KEY**

### PART 1: MOLECULAR BIOLOGY BASICS

*Instructor Note: PART 1 is modified from the activity "POGIL Gene Expression – Transcription", POGIL$^{TM}$ Activities for AP* Biology. We recommend using whatever portions of the activity are appropriate for your course in providing a review of the central dogma.*

### PART 2: BACTERIAL GENOMIC ORGANIZATION

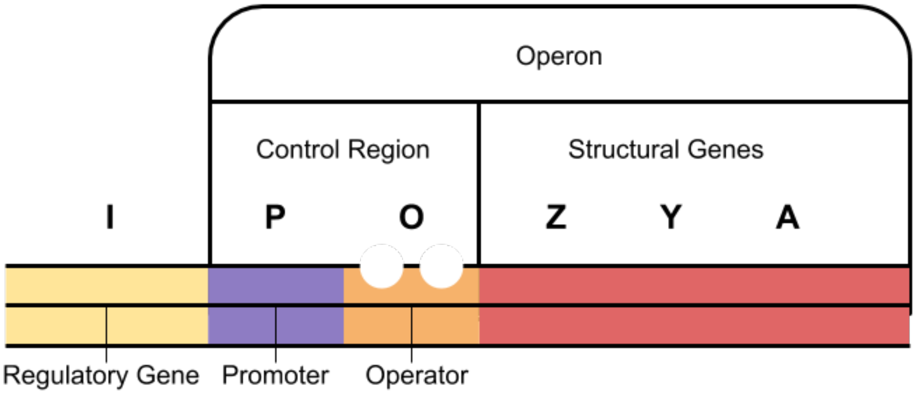


1. Refer to the image above to answer the following questions.

   a. How many genes are contained in this operon?
      *three*
   b. How many promoters control the expression of this operon?
      *one*
   c. How many mRNAs would be transcribed from this operon?
      *one*
   d. How many polypeptides would be produced?
      *three*
   e. How many start codons, and how many stop codons would be present in this operon?
      *three*
   f. How many ribosome binding sites (RBS) would be present in this operon?
      *three*

### PART 3: BACTERIAL TRANSCRIPTION

1. Where in the eukaryotic cell does transcription take place? *in the nucleus*
2. Where in the prokaryotic cell does transcription take place? *in the cytoplasm*
3. List the components that would be included a bacterial **promoter** region. *-35 element, -10 element*

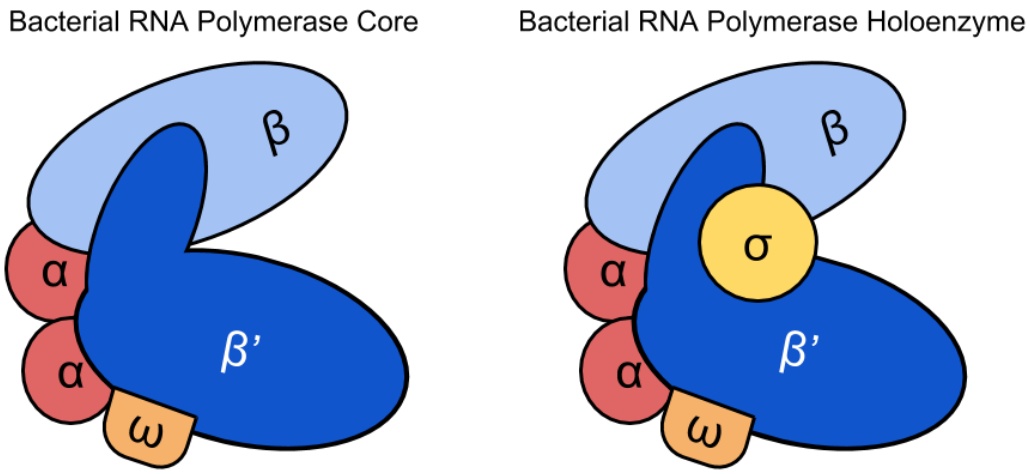


4. The image above shows the bacterial RNA polymerase core and holoenzyme complexes. What additional subunit is present in the holoenzyme that is not part of the core?
   *Sigma subunit*
5. Which subunit(s) do you think would bind each of the promoter regions that you listed in Q3?
   *Sigma subunit*

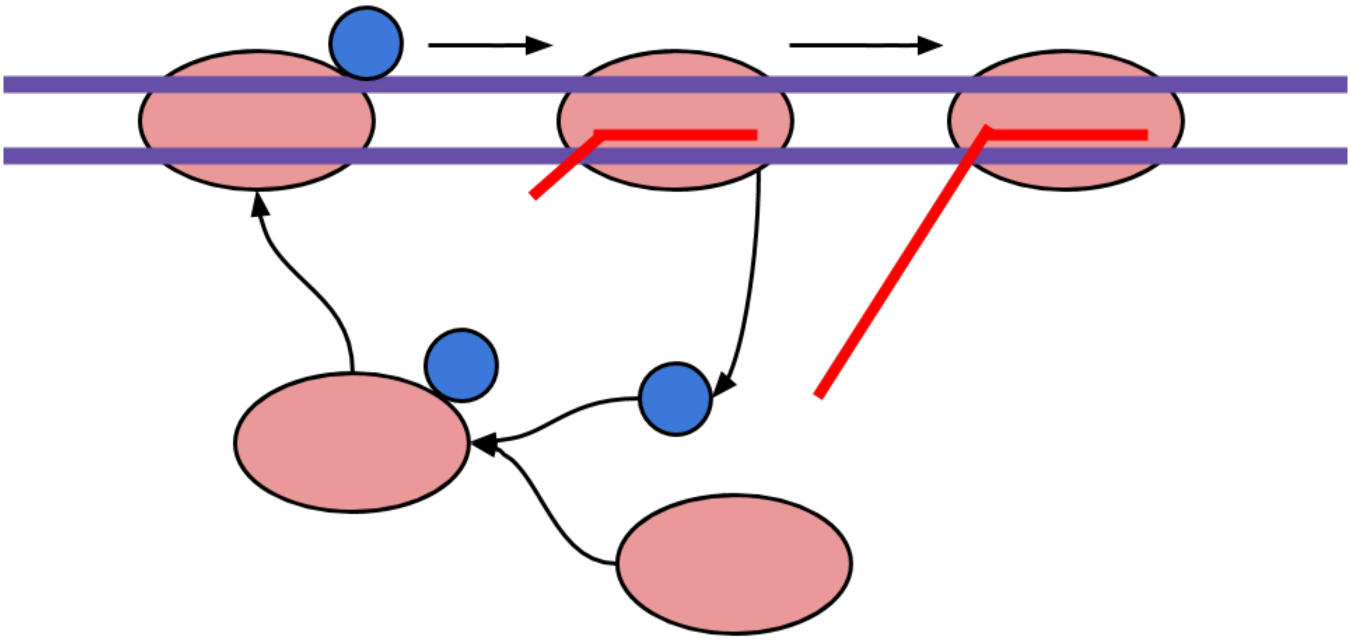

6. The image above shows a cycle that occurs during transcription initiation and elongation.

   a. What do you think the blue ball represents?
      *Sigma subunit*
   b. What do you think the red oval represents?
      *RNA polymerase core*
   c. What do you think the red line represents?
      *A nascent transcript*

d. Describe what is happening in this image.
*This image depicts the sigma cycle. Student descriptions will differ, but generally, they should note that the holoenzyme binds to the DNA and begins to move along the DNA as it transcribes. After transcription is initiated, the sigma subunit is released. The RNA polymerase core continues transcribing the gene, and the sigma subunit can bind to another RNA polymerase core molecule to begin another round of transcription of this gene, or transcription of another gene.*

#### PART 4: BACTERIAL TRANSLATION

*Instructor Note: PART 4 is modified from the activity "POGIL Gene Expression – Translation", POGIL™ Activities for AP* Biology. We recommend using whatever portions of the activity are appropriate for your course in providing a review of translation.*

**STOP**

**COMPLETE PART 5 WITH THE MODEL PROVIDED NOW.**

#### PART 5: REGULATION OF BACTERIAL GENE EXPRESSION

1. Examine the red/white model.
   a. What do you think the red portion represents?
   *RNA polymerase core*
   b. What do you think the white portion represents?
   *Sigma subunit*
2. Examine the gray DNA operon model. *Note- this model is only showing you the top strand (5'>3') for simplicity.

   a. How many promoters control the expression of this operon?
   *One*
   b. Place the "feet" of the red/white model into the track on the left most side of the DNA. Slowly slide it towards the right, along the sequence until you arrive at the -35 and -10 regions.
   *Students should be able to locate the -35 and -10 sites as the motor will vibrate when the polymerase contacts these regions of the DNA box.*
   c. Underline the -35 and -10 consensus sequences using the black sharpie. (you can refer back to previous lecture notes!)
   *Students should underline TTTACA (-35 sequence) and TTATGTT (-10 sequence). Of note, the -35 and -10 element sequences found in the lac operon promoter differ from the consensus sequences of TTGACA and TATAAT, respectively.*

   d. Mark the approximate location +1 with an *
   *Students should add an * to indicate the +1 site. After underlining and adding the asterisk, their sequence label should read:*

CCAGGCTTTACACTTTCTGCTTCCTTCGTTATGTTGTCGGA* ATTGTGAGCGGATAACAATT

4. Examine the gray DNA operon model. *Note- this model is only showing you the top strand (5'>3') for simplicity.

   a. List the genes that are contained in this operon.
   *lacZ, lacY, and lacA*
   b. How many mRNAs would be transcribed from this operon?
   *One*
   c. How many polypeptides would be produced?
   *Three*
   d. With your black sharpie, mark the location where you would find the START and STOP codon(s).
   *Students should place a start codon at the beginning of each gene, and a stop codon at the end of each gene (three start codons and three stop codons).*
   e. With your other color sharpie, mark the location where you would find the RBS(s).
   *Students should place an RBS upstream of each of the three start codons.*

8. Using the foam "protein", determine where on your DNA operon model this protein will bind.

   a. Underline this sequence (does not need to be precise) with your colored sharpie.
   *Students should underline sequence where the magnet has been glued into the DNA operon box. The lacI binding site begins at the +1 site, but depending on the preciseness of the label placement and magnet placement, students might place the binding site near, but not exactly at, the +1 site.*
   b. What protein does the foam protein represent?
   *lacI (repressor)*

9. Allow the pink marble to "interact" with the foam protein.

   a. How does the pink molecule interacting with the foam protein impact the red/white model from performing its function?
   *When the pink molecule (marble) is not present, the foam protein binds to the DNA and prevents the red/white model (polymerase) from binding to the promoter region. When the pink molecule (marble) binds to the foam protein, the protein can no longer bind to the DNA sequence. This allows the polymerase to access the promoter.*

   b. What molecule does the pink molecule represent?
   *Allolactose*

10. If the pink marble is not present, is the transcription of this operon turned on or off?
    *Off*
11. If the pink marble is present, is the transcription of this operon turned on or off?
    *On*

#### PART 6: REGULATION OF BACTERIAL GENE EXPRESSION

Recall that glucose is used to generate energy in the form of ATP through glycolysis, and in some organisms, through oxidative phosphorylation. It is the preferred carbon source for *E. coli*.

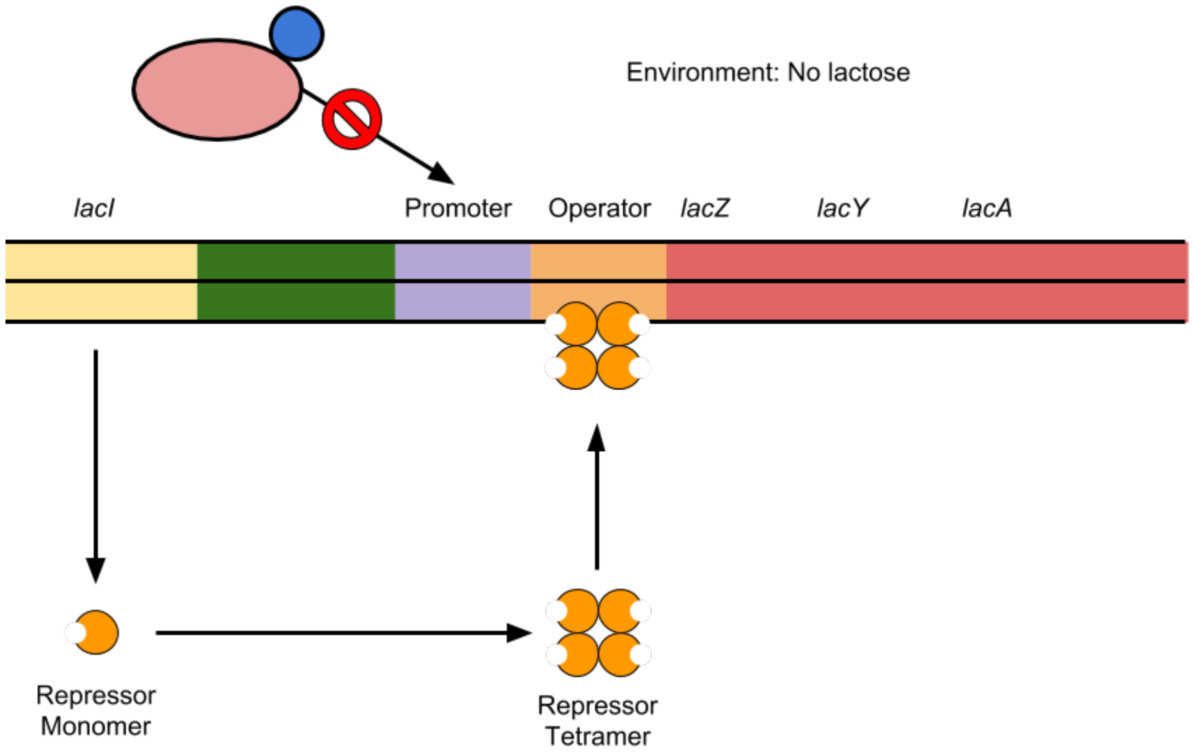


1. Refer to the image above to answer the following questions.

   a. What is encoded by the *lacI* gene?
      *Repressor monomer*
   b. Is the *lacI* gene part of the lac operon?
      *No*
   c. What happens to the protein encoded by the *lacI* gene in the absence of lactose?
      *It forms a tetramer and binds to DNA*
   d. To what site in the DNA does the protein bind?
      *The lac operator*
   e. How does this affect the transcription of the genes in the *lac* operon?
      *This inhibits transcription by preventing the RNA polymerase holoenzyme from binding the promoter.*

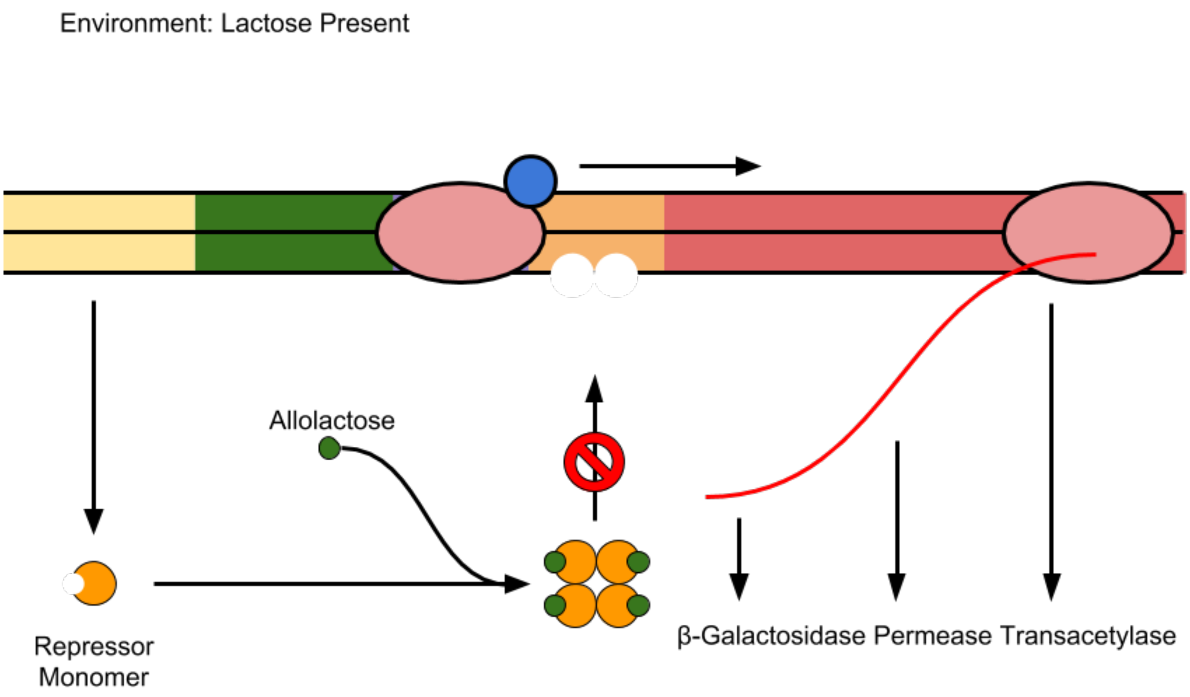


2. Refer to the image above to answer the following questions. Note, when lactose is present in large enough quantities, some of this lactose will be converted to allolactose.

   a. What molecule is considered the "inducer" in this system?
   *Allolactose*
   b. What molecule or molecules are bound by the inducer?
   *The repressor tetramer*
   c. How does that affect the function of these molecules?
   *It prevents the repressor tetramer from binding to the lac operator*
   d. How does this affect the transcription of the genes in the *lac* operon?
   *The RNA polymerase holoenzyme is not blocked from binding to the promoter, and transcription can occur.*

3. Considering both images together, how would you expect transcriptional activity at the *lac* operon to change if you added large quantities of lactose to an *E. coli* culture?
*If large quantities of lactose are added to an E. coli culture, transcription of the lac operon will be induced.*

4. How would you expect transcriptional activity at the *lac* operon to change if you removed lactose from an *E. coli* culture?
*If you remove lactose from an E. coli culture, lac operon transcription will be inhibited.*

5. Do you think that glucose levels should also regulate *lac* operon transcription? If so, how? That is, should high levels of glucose increase or decrease *lac* operon transcription?

*Yes: if high levels of glucose are present, it is not necessary for the cell to use lactose, even if large quantities of lactose are present. High levels of glucose should decrease lac operon transcription.*

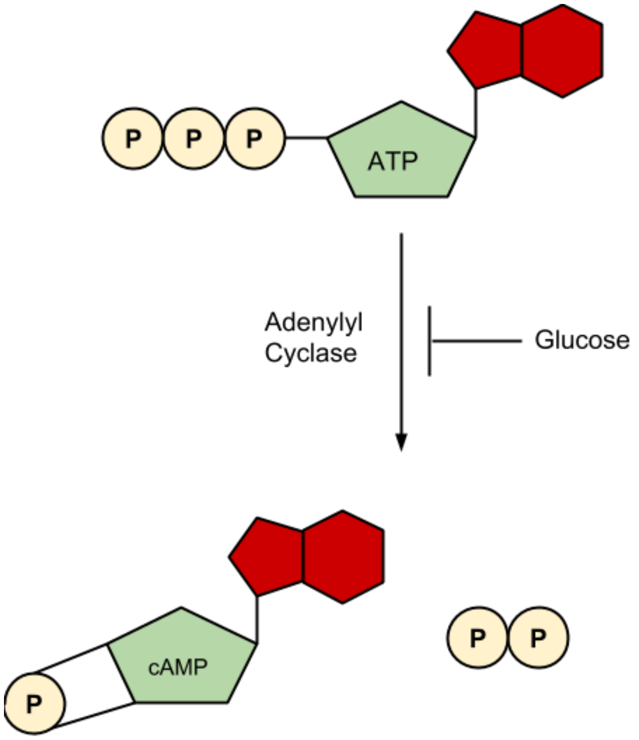


6. Describe the chemical reaction above in one or two sentences.
   *Adenylyl cyclase converts ATP into cAMP. This reaction is inhibited by glucose.*

7. When glucose levels in the cell decrease, would you expect an increase or decrease in cAMP levels? Why?
   *When glucose levels decrease, less glucose will be present to inhibit the action of adenylyl cyclase. This will result in increased levels of cAMP.*

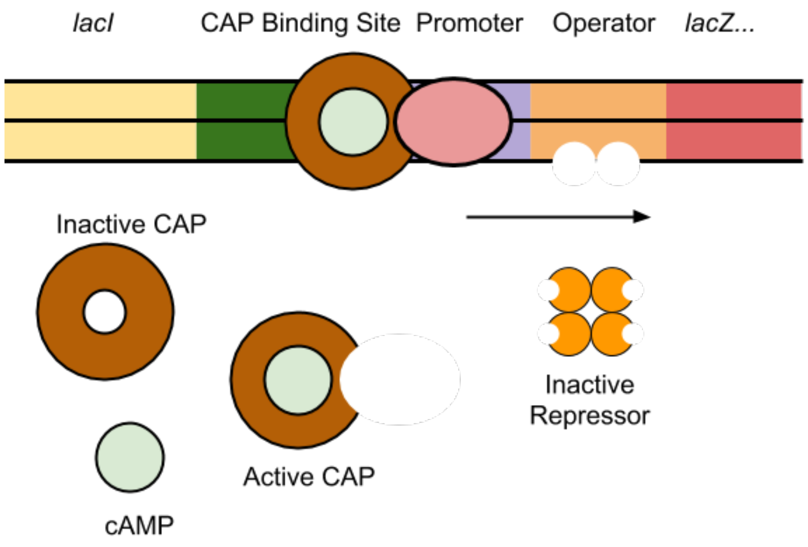


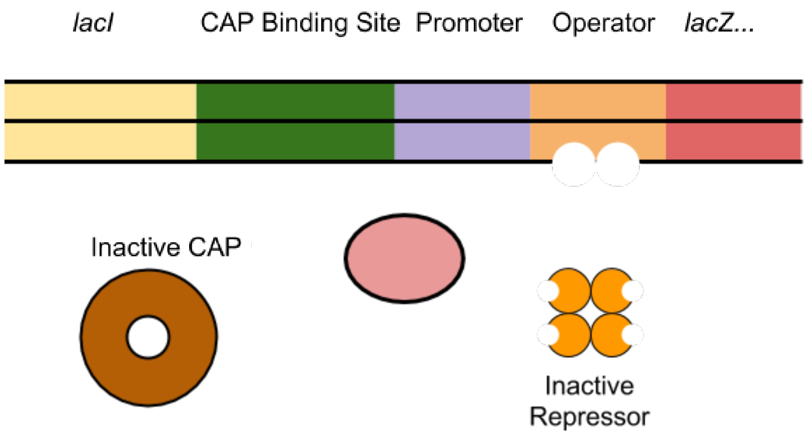


8. Refer to the image above to answer the following questions. In the images above, we have added some additional detail.

   In the top image, lactose is present, and glucose is absent.

   a. Where is the repressor? Why?
      *The repressor is not bound to the DNA. Because lactose is present, the repressor tetramer binds allolactose, which prevents it from binding to the lac operator.*
   b. Are cAMP levels high or low? Why?
      *cAMP levels are high because glucose is absent, and there is decreased inhibition of adenylyl cyclase.*
   c. What protein is cAMP interacting with?
      *cAMP is interacting with CAP.*
   d. What does this protein do when it interacts with cAMP?
      *CAP becomes activated and binds to the CAP binding site in the DNA.*

e. How does this affect the transcription of the genes in the *lac* operon?
   *This activates transcription of the lac operon.*

In the bottom image, lactose is present, and glucose is present.

f. Where is the repressor? Why?
   *The repressor is not bound to the DNA. Because lactose is present, the repressor tetramer binds allolactose, which prevents it from binding to the lac operator.*
g. Are cAMP levels high or low? Why?
   *cAMP levels are low because glucose is present, and glucose inhibits adenylyl cyclase.*
h. What happens to CAP?
   *It remains inactive.*
i. Are the genes in the *lac* operon transcribed?
   *No.*

#### PART 7: APPLYING THIS INFORMATION

1. What conditions are necessary for transcription of the *lac* operon to occur?
   *Transcription of the lac operon occurs when lactose levels are high and glucose levels are low.*

2. Would you say that lactose directly or indirectly regulates the *lac* operon?
   *Lactose indirectly regulates the lac operon by regulating the ability of the lacI protein to bind to the lac operator.*

3. Would you say that glucose directly or indirectly regulates the *lac* operon?
   *Glucose indirectly regulates the lac operon by regulating the ability of adenylyl cyclase to convert ATP to cAMP. (cAMP, in turn, is also an indirect regulator, as it binds to and activates CAP, a direct regulator of lac operon transcription.)*

4. Come up with an example of a mutation in one component of the *lac* operon, and answer the following questions.

   a. What component is mutated, and what is the mutation?

   b. In your altered version of the lac operon, the presence of lactose and absence of glucose, is the lac operon transcribed?

   c. Explain, or draw, why this is the case.

*Examples of possible answers*

| ***Possible mutations*** | ***Is repressor bound to DNA?*** | ***Is that lac operon expressed?*** |
|---|---|---|
| *mutation in operator prevents repressor from binding* | *no* | *yes* |
| *mutation in repressor prevents it from binding allolactose* | *yes* | *no* |

| | | |
|---|---|---|
| *mutation in operator keeps repressor bound* | *yes* | *no* |

**Appendix 5: Pre and post assessment questions KEY**

**Pre-assessment:** The pre-assessment consisted of questions from the case study entitled "Cow of the Future Genetically Engineering a Microbe to Reduce Bovine Methane Emissions" (Stewart et al, 2014).

**Post-assessment:** The following exam questions were used for post-assessment. **Correct answers are bolded and in red.**

The following is a diagram of the *ara* operon in *E. coli*. This operon encodes the enzymes needed to utilize the sugar arabinose.

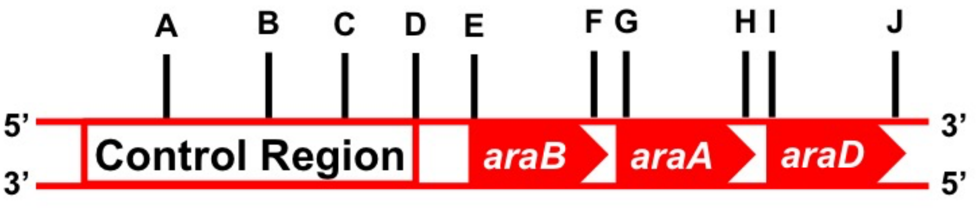


1. How many genes are contained in this operon?
   Select one:
   - 1
   - 2
   - **3**
   - 4
   - 5
   - 6

2. How many mRNAs would be transcribed from this operon?
   Select one:
   - **1**
   - 2
   - 3
   - 4
   - 5
   - 6

3. How many polypeptides would be produced?
   Select one:
   - 1
   - 2
   - **3**
   - 4
   - 5

6

4. How many promoters control expression of this operon?
Select one:

**1**

2

3

4

5

6

5. How many start codons would be present in this operon?
Select one:

1

2

**3**

4

5

6

6. How many stop codons would be present in this operon?
Select one:

1

2

**3**

4

5

6

**Appendix 6: Blank student worksheet**

#### BACTERIAL GENOMIC ORGANIZATION

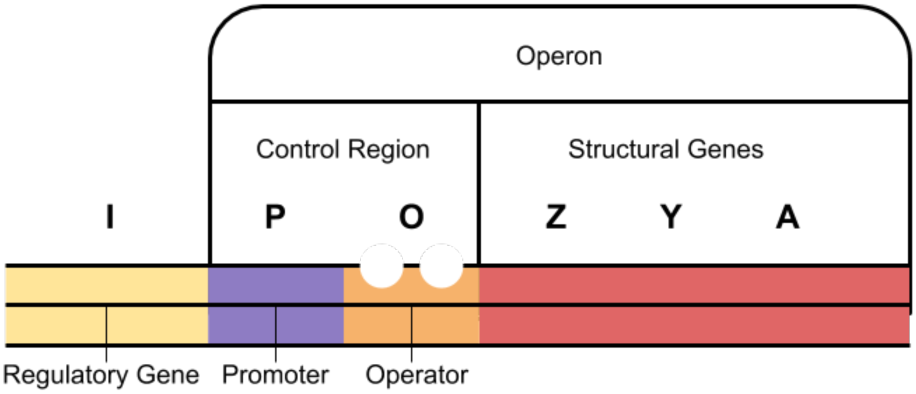


1. Refer to the image above to answer the following questions.

   a. How many genes are contained in this operon?

   b. How many promoters control the expression of this operon?

   c. How many mRNAs would be transcribed from this operon?

   d. How many polypeptides would be produced?

   e. How many start codons, and how many stop codons would be present in this operon?

   f. How many ribosome binding sites (RBS) would be present in this operon?

#### BACTERIAL TRANSCRIPTION

1. Where in the eukaryotic cell does transcription take place?
2. Where in the prokaryotic the cell does transcription take place?
3. List the components that would be included in a bacterial **promoter** region.

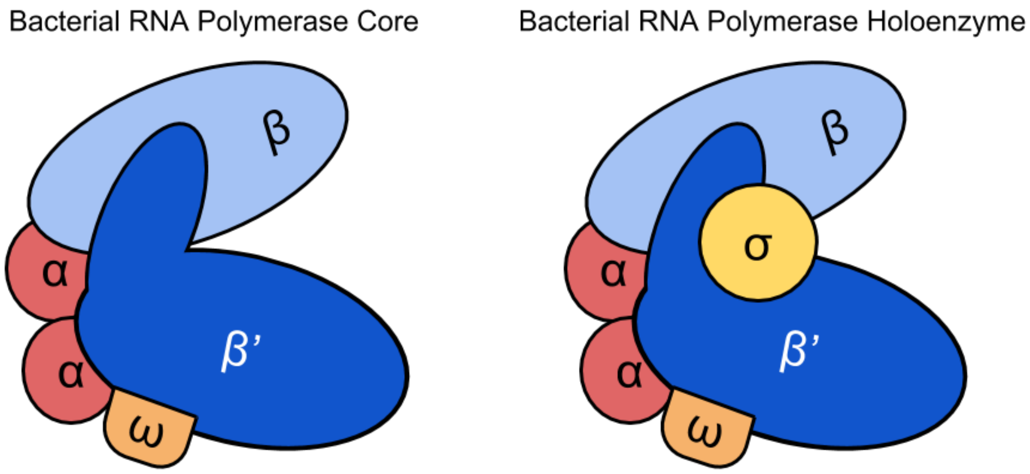


4. The image above shows the bacterial RNA polymerase core and holoenzyme complexes. What additional subunit is present in the holoenzyme that is not part of the core?

5. Which subunit(s) do you think would bind each of the promoter regions that you listed in Q3?

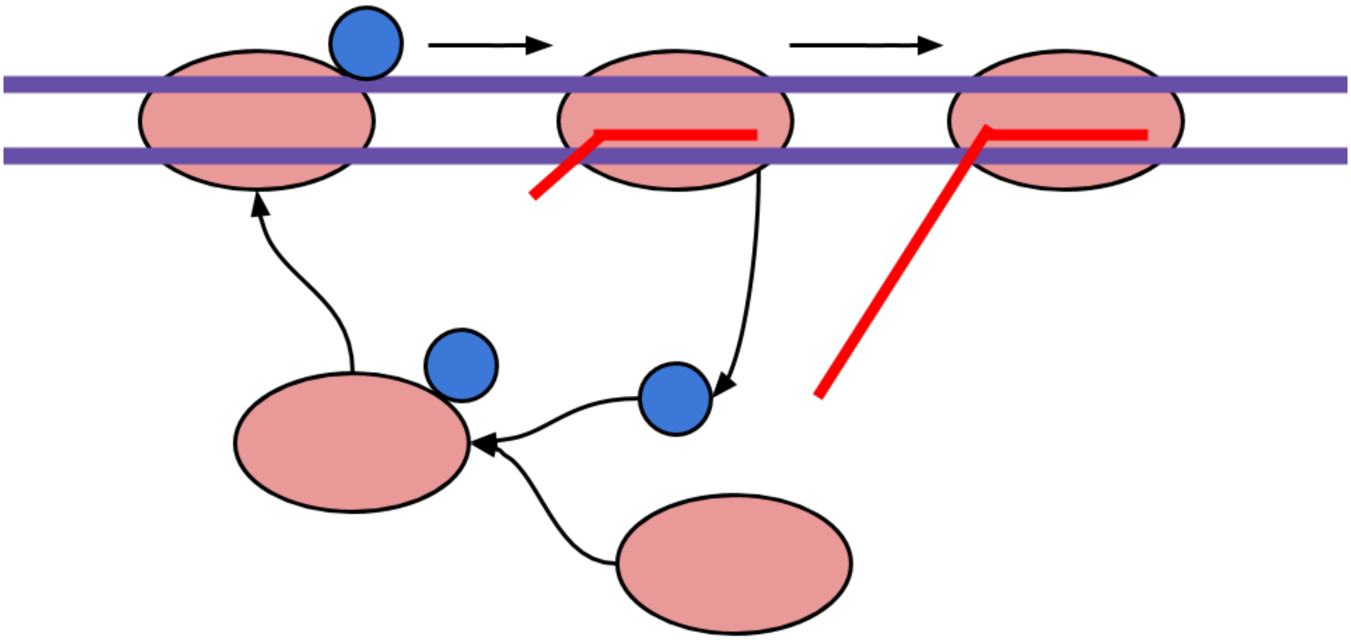

6. The image above shows a cycle that occurs during transcription initiation and elongation.

   a. What do you think the blue ball represents?

   b. What do you think the red oval represents?

   c. What do you think the red line represents?

d. Describe what is happening in this image.

**STOP**

**COMPLETE PART 5 WITH THE MODEL PROVIDED NOW.**

#### REGULATION OF BACTERIAL GENE EXPRESSION

1. Examine the red/white model.
   a. What do you think the red portion represents?

   b. What do you think the white portion represents?

2. Examine the gray DNA operon model. *Note- this model is only showing you the top strand (5'>3') for simplicity.

   a. How many promoters control the expression of this operon?

   b. Place the "feet" of the red/white model into the track on the left most side of the DNA. Slowly slide it towards the right, along the sequence until you arrive at the -35 and -10 regions.

   c. Underline the -35 and -10 consensus sequences using the black sharpie.

   d. Mark the approximate location +1 with an *

3. Examine the gray DNA operon model. *Note- this model is only showing you the top strand (5'>3') for simplicity.

   a. List the genes that are contained in this operon.

   b. How many mRNAs would be transcribed from this operon?

   c. How many polypeptides would be produced?

   d. With your black sharpie, mark the location where you would find the START and STOP codon(s).

   e. With your other color sharpie, mark the location where you would find the RBS(s).

4. Using the foam "protein", determine where on your DNA operon model this protein will bind.

a. Underline this sequence (does not need to be precise) with your color sharpie.

b. What protein does the foam protein represent?

5. Allow the pink marble to "interact" with the foam protein.

a. How does the pink molecule interacting with the foam protein impact the red/white model from performing its function?

b. What molecule does the pink molecule represent?

6. If the pink marble is not present, is the transcription of this operon turned on or off?

7. If the pink marble is present, is the transcription of this operon turned on or off?

#### REGULATION OF BACTERIAL GENE EXPRESSION

Recall that glucose is used to generate energy in the form of ATP through glycolysis, and in some organisms, through oxidative phosphorylation. It is the preferred carbon source for *E. coli*.

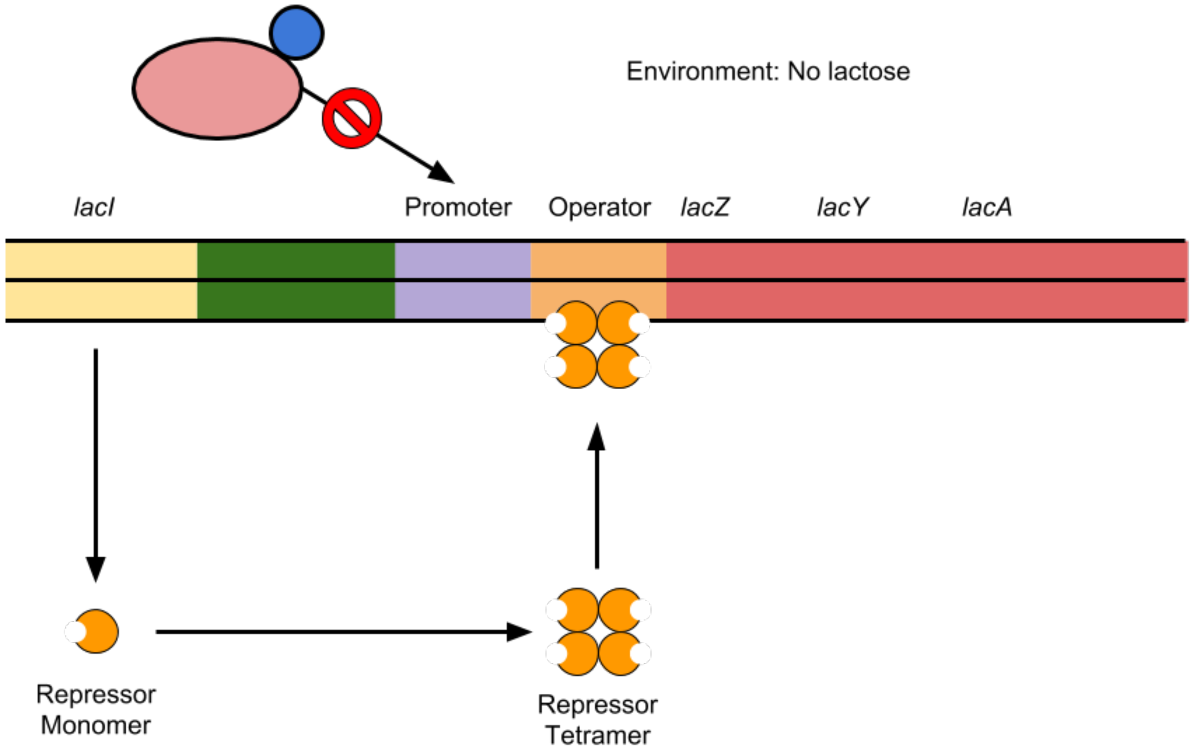


1. Refer to the image above to answer the following questions.

a. What is encoded by the *lacI* gene?

b. Is the *lacI* gene part of the lac operon?

c. What happens to the protein encoded by the *lacI* gene in the absence of lactose?

d. To what site in the DNA does the protein bind?

e. How does this affect the transcription of the genes in the *lac* operon?

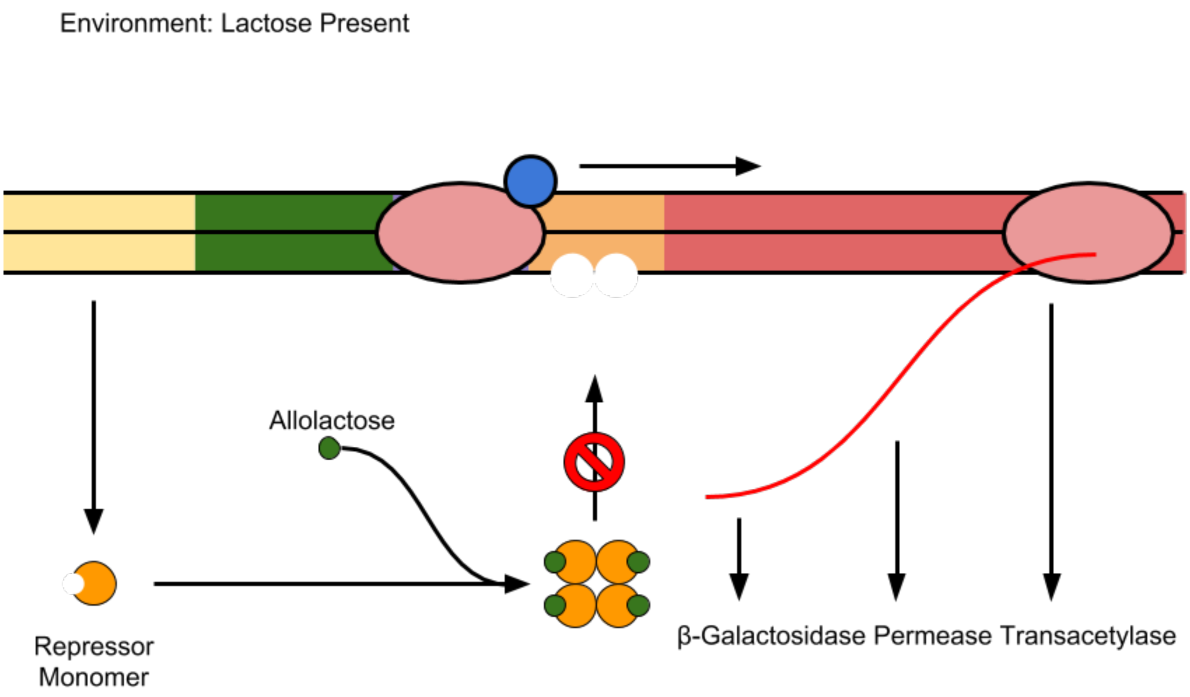


2. Refer to the image above to answer the following questions. Note, when lactose is present in large enough quantities, some of this lactose will be converted to allolactose.

   a. What molecule is considered the "inducer" in this system?

   b. What molecule or molecules are bound by the inducer?

   c. How does that affect the function of these molecules?

   d. How does this affect the transcription of the genes in the *lac* operon?

3. Considering both images together, how would you expect transcriptional activity at the *lac* operon to change if you added large quantities of lactose to an *E. coli* culture?

4. How would you expect transcriptional activity at the *lac* operon to change if you removed lactose from an *E. coli* culture?

5. Do you think that glucose levels should also regulate *lac* operon transcription? If so, how? That is, should high levels of glucose increase or decrease *lac* operon transcription?

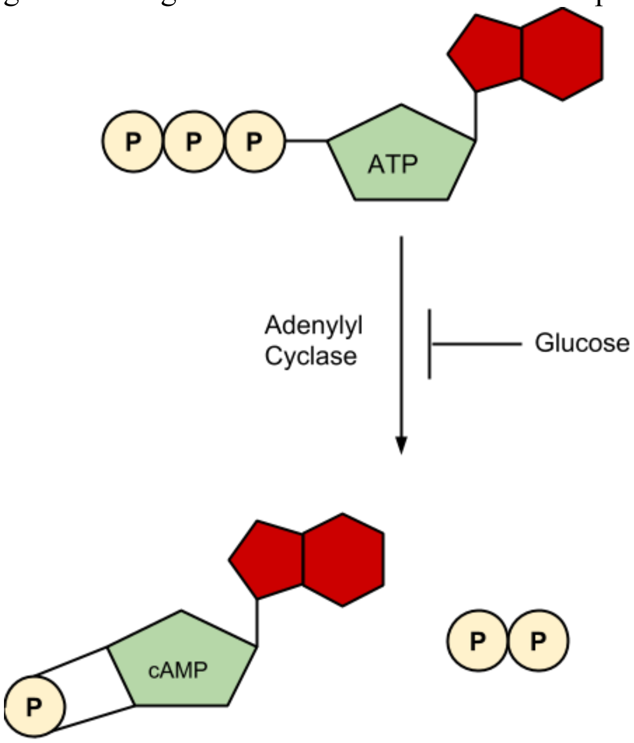


6. Describe the chemical reaction to the left in one or two sentences.

7. When glucose levels in the cell decrease, would you expect an increase or decrease in cAMP levels? Why?

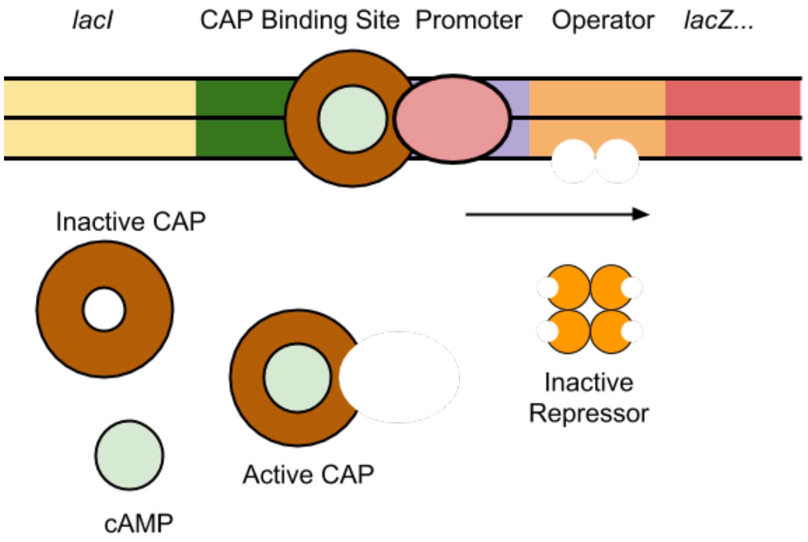


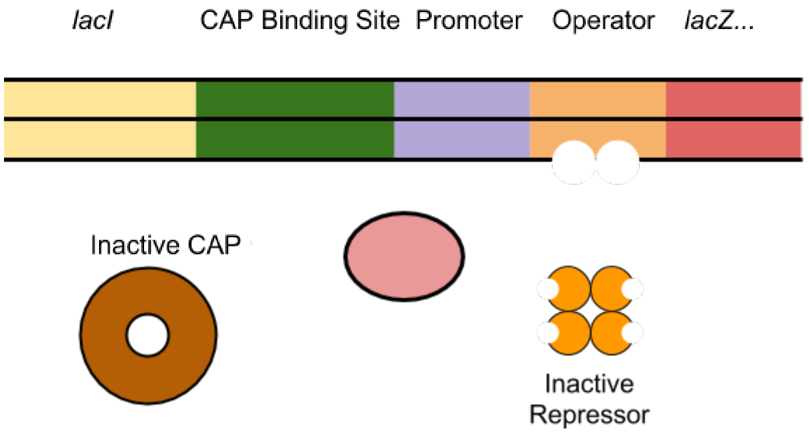


8. Refer to the image above to answer the following questions. In the images above, we have added some additional detail.

   In the top image, lactose is present, and glucose is absent.

   a. Where is the repressor? Why?

   b. Are cAMP levels high or low? Why?

   c. What protein is cAMP interacting with?

   d. What does this protein do when it interacts with cAMP?

e. How does this affect the transcription of the genes in the *lac* operon?

In the bottom image, lactose is present, and glucose is present.

f. Where is the repressor? Why?

g. Are cAMP levels high or low? Why?

h. What happens to CAP?

i. Are the genes in the *lac* operon transcribed?

#### APPLYING THIS INFORMATION

1. What conditions are necessary for transcription of the *lac* operon to occur?

2. Would you say that lactose directly or indirectly regulates the *lac* operon?

3. Would you say that glucose directly or indirectly regulates the *lac* operon?

4. Come up with an example a mutation in one component of the *lac* operon, and answer the following questions.

    a. What component is mutated, and what is the mutation?

    b. In your altered version of the *lac* operon, the presence of lactose and absence of glucose, if the *lac* operon transcribed?

    c. Explain, or draw, why this is the case.
